# Enhancers with cooperative Notch binding sites are more resistant to regulation by the Hairless co-repressor

**DOI:** 10.1101/2020.07.12.199422

**Authors:** Yi Kuang, Anna Pyo, Natanel Eafergan, Brittany Cain, Lisa M. Gutzwiller, Ofri Axelrod, Ellen K. Gagliani, Matthew T. Weirauch, Raphael Kopan, Rhett A. Kovall, David Sprinzak, Brian Gebelein

## Abstract

Notch signaling controls many developmental processes by regulating gene expression. Notch-dependent enhancers recruit activation complexes consisting of the Notch intracellular domain, the <u>C</u>bf/<u>S</u>u(H)/<u>L</u>ag1 (CSL) transcription factor (TF), and the Mastermind co-factor via two types of DNA sites: monomeric CSL sites and cooperative dimer sites called <u>S</u>u(H) <u>p</u>aired <u>s</u>ites (SPS). Intriguingly, the CSL TF can also bind co-repressors to negatively regulate transcription via these same sites. Here, we tested how enhancers with monomeric CSL sites versus dimeric SPSs bind *Drosophila* Su(H) complexes *in vitro* and mediate transcriptional outcomes *in vivo*. Our findings reveal that while the Su(H)/Hairless co-repressor complex similarly binds SPS and CSL sites in an additive manner, the Notch activation complex binds SPSs, but not CSL sites, in a cooperative manner. Moreover, transgenic reporters with SPSs mediate stronger, more consistent transcription and are more resistant to increased Hairless co-repressor expression compared to reporters with the same number of CSL sites. These findings support a model in which SPS containing enhancers preferentially recruit cooperative Notch activation complexes over Hairless repression complexes to ensure consistent target gene activation.

## Introduction

Notch signaling is a highly conserved cell-to-cell communication pathway that conveys information required for proper cellular decisions in many tissues and organs. During embryonic development, the Notch signaling pathway is used to specify distinct cell fates and thereby plays crucial roles during organogenesis including vasculogenesis (Siekmann and Lawson, 2007), hematopoiesis (Carlesso et al., 1999), neurogenesis (Ahmad et al., 1995; Xu et al., 1990), and cardiac development (McCright et al., 2001; Park et al., 1998; Rones et al., 2000). Additionally, Notch regulates tissue homeostasis, including epidermal differentiation and maintenance (Heitzler and Simpson, 1991), lymphocyte differentiation (Robson MacDonald et al., 2001), muscle and bone regeneration (Fukushima et al., 2017; Hilton et al., 2008; Koch et al., 2013), and angiogenesis (Siekmann and Lawson, 2007). Intriguingly, Notch regulates these diverse processes using a common molecular cascade that is initiated through a ligand (Delta/Serrate/Jagged)-receptor (Notch) interaction that triggers the cleavage and release of the Notch intracellular domain (NICD) into the cytoplasm. NICD subsequently translocates into the nucleus and forms a ternary complex with the Cbf1/Su(H)/Lag1 (CSL) transcription factor (which is also commonly called RBPJ in mammals) and the Mastermind (Mam) adapter protein. The NICD/CSL/Mam (NCM) complex recruits the p300 co-activator to activate the expression of Notch target genes required for proper cellular outcomes (Kopan and Ilagan, 2009; Kovall et al., 2017).

Since NICD and Mam do not directly bind DNA, the targeting of the NCM complex to specific genomic loci is determined by the CSL transcription factor (TF). Both *in vitro* and *in vivo* DNA binding assays show that the CSL TFs from *C elegans, Drosophila*, and vertebrates bind highly similar DNA sequences (i.e. ^T^/_C_GTG^G^/_A_GAA), and its interactions with Notch and Mam do not alter CSL DNA binding specificity (del Bianco et al., 2010; Castel et al., 2013; Christensen et al., 1996; Fortini and Artavanis-Tsakonas, 1994; Friedmann and Kovall, 2010; Tamura et al., 1995; Tun et al., 1994). Interestingly, studies in flies and mammals found that a subset of Notch target genes contain enhancers with two binding sites spaced 15 to 17bp apart and oriented in a head-to-head manner (Bailey and Posakony, 1995; Nellesen et al., 1999; Severson et al., 2017). Subsequent biochemical and structural studies revealed that such sites, which have been named <u>S</u>u(H) <u>p</u>aired <u>s</u>ites or <u>s</u>equence <u>p</u>aired <u>s</u>ites (SPSs), mediate cooperative NCM binding due to the dimerization between two adjacent NICD molecules (Arnett et al., 2010; Nam et al., 2007).

SPSs are present in a substantial fraction of Notch-dependent enhancers in the genome. In human CUTLL1 T-cell acute lymphoblastic leukemia (T-ALL) cell line, 36% (38 of 107) of the high confident Notch targets are dimer-dependent (Severson et al., 2017), and SPS-containing enhancers were found to be crucial for the maturation of both normal T-cells and the progression of T-ALL (Liu et al., 2010; Yashiro-Ohtani et al., 2014). A genome-wide NICD complementation assay revealed that mouse mK4 kidney cells have as many as 2,500 Notch dimer-dependent loci (Hass et al., 2015). Moreover, reporter assays and/or RT-PCR assays have tested the function of a small subset of these SPS containing enhancers and found that SPSs are typically required for optimal transcriptional responses (Arnett et al., 2010; Hass et al., 2015; Liu et al., 2010; Nam et al., 2007). A recent study also found that while mice with Notch1 and Notch2 point mutations that abolish cooperative binding to SPSs develop normally under ideal laboratory conditions, stressing the animals either genetically or with parasites can result in profound defects in gastrointestinal, cardiovascular, and immune systems (Kobia et al., 2020). Collectively, these studies revealed that a large number of Notch-dependent target genes contain SPSs, and that the regulation of dimer-dependent Notch target genes contributes to animal development and homeostasis.

In addition to mediating Notch induced gene expression, the CSL TF can use the same DNA binding sites to repress transcription by recruiting co-repressor proteins. The *Drosophila* CSL transcription factor Su(H) binds to the co-repressor Hairless (H) protein, which recruits either the Groucho (Gro) or the C-terminal binding protein (Ctbp) co-repressors (Barolo et al., 2002; Morel et al., 2001). The mammalian CSL transcription factor RBPJ interacts with several transcriptional repressors including the SHARP/Mint protein and Fhl1C/KyoT2 (Kovall and Blacklow, 2010). Once bound to DNA, these co-repressor complexes recruit additional proteins that can mediate transcriptional repression by modifying chromatin. Importantly, recent structural analysis of the fly and mammalian co-repressors bound to CSL and DNA revealed that co-repressors interact with the CSL TF in a competitive manner with the NICD/Mam activation complex (Maier et al., 2011; Yuan et al., 2016, 2019). Moreover, genetic studies revealed that the ratio of the co-activator to co-repressor complex is critical for proper Notch-mediated cellular decisions, as lowering the gene dose of the *Hairless* co-repressor can suppress *Notch* haploinsufficiency phenotypes in *Drosophila* (Price et al., 1997). In total, these data support a model whereby the Notch activation complex directly competes for genomic binding sites with the CSL/co-repressor complex to regulate target gene expression.

Recent studies have begun to focus on defining whether Notch regulated enhancers with SPSs convey distinct transcriptional responses from CSL monomeric sites. For example, the *E(spl)* genes, many of which contain SPSs, were found to be among the first to respond after a short pulse of Notch activation in *Drosophila* DmD8 cells (Housden et al., 2013), consistent with SPS-containing enhancers responding quickly to low levels of Notch activation. However, subsequent live imaging studies comparing the activities of enhancers with SPS *versus* CSL sites revealed that the presence of SPSs did not significantly alter the sensitivity to NICD but instead enhanced transcriptional burst size (Falo-Sanjuan et al., 2018). It should be noted, however, that these studies have largely focused on how the Notch activation complex cooperatively binds to and impacts the regulation of SPS containing enhancers, whereas less is known about whether and how the SPS versus monomeric CSL sites differentially recruit the CSL/co-repressor complexes. Thus, it remains unclear how the levels of the co-repressors impact Notch regulated enhancers that contain cooperative SPS sites versus independent CSL sites.

Comparing Notch-mediated transcriptional responses of endogenous enhancers with SPS and CSL sites is complicated by several inherent properties of endogenous enhancers. First, most Notch-regulated SPS-containing enhancers also have variable numbers of independent monomer CSL sites. Second, endogenous enhancers contain distinct combinations of additional TF binding sites that can significantly alter transcriptional output. Third, each endogenous enhancer is embedded in its own unique chromosomal environment, which can further impact the ability of Notch transcription complexes to regulate gene expression. In this study, we circumvented these confounders by integrating transgenic reporters containing either synthetic SPS or CSL enhancers to focus our investigation on how the architecture of Su(H) binding sites impacts Notch transcriptional output in *Drosophila*. We complemented these studies using *in vitro* DNA binding assays to assess how SPS versus CSL sites impact the binding of the NCM versus CSL/co-repressor complexes. Altogether, our data reveal that Notch regulated enhancers containing cooperative SPSs are more resistant to the Hairless co-repressor protein than enhancers with independent CSL sites. Integrating this study with previously published data provides new insights into how the architecture of CSL binding sites affect transcriptional output by both modulating transcriptional dynamics and by competing with the co-repressors that limit transcriptional activation.

## Results

### Activating but not repressing Su(H) complexes cooperatively bind SPS sites in vitro

To study the ability of CSL vs SPS sites to bind activating (NICD/CSL/MAM, NCM) and repressing (CSL/Hairless) complexes that regulate gene expression in *Drosophila*, we designed synthetic CSL and SPS enhancers for *in vivo* transgenic reporter assays. To isolate the Su(H) (the *Drosophila* CSL TF) binding sites from additional potential transcriptional inputs, the intervening sequences were selected to exclude other known TFBSs by randomly generating thousands of sequence variants and scoring each using a TF binding motif database (CIS-BP, http://cisbp.ccbr.utoronto.ca) (Weirauch et al., 2014). High-affinity Notch monomer sites (CSL) were designed in a head-to-tail fashion with sufficient spacing (17 bp) to permit independent binding of NCM complexes, whereas the same high-affinity 8-mer sequence was oriented in a head-to-head manner 15 bp apart to generate an SPS site capable of cooperative NCM binding (**Fig 1A**).

**Figure 1.**
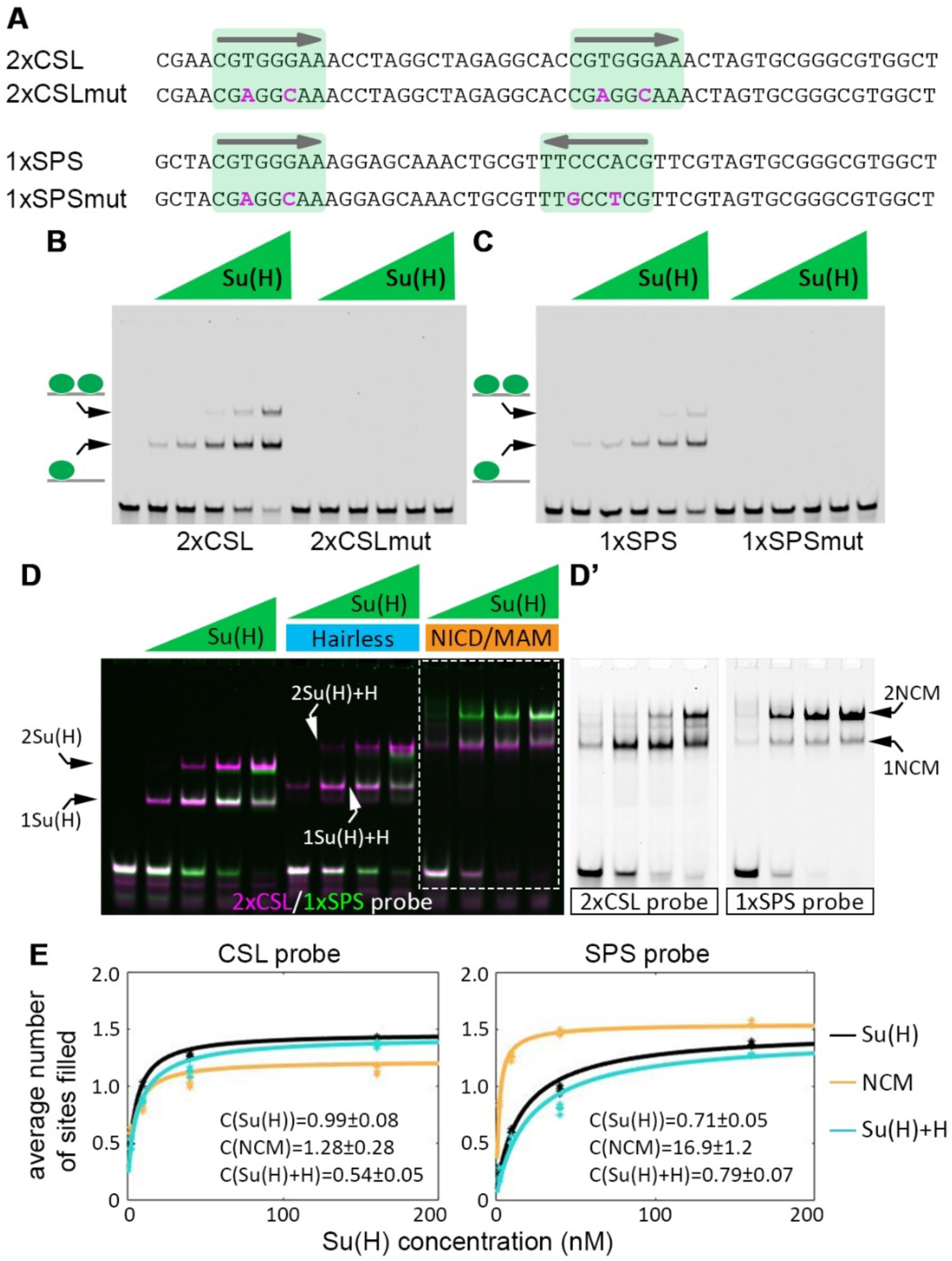
The Notch-CSL-Mastermind (NCM) complex binds the SPS sequence cooperatively *in vitro*. **A**. Sequences of the 2xCSL and 1xSPS probes, which both contain two of the same Su(H) binding site (CGTGGGAA, highlighted in green) that only differ in orientation and spacing. The specific mutations introduced into the Su(H) binding sites are noted in magenta text. **B-C**. EMSAs reveal binding of purified *Drosophila* Su(H) to the wild type, but not the mutated, 2xCSL (B) and 1xSPS (C) probes. Su(H) concentration increases from 0.94nM to 15nM in 2-fold steps. **D**. EMSA reveals binding of the indicated purified *Drosophila* proteins on 2xCSL (magenta) or 1xSPS (green) probes. Note, Su(H) alone and the Su(H)/H co-repressor complex bind the 2xCSL and 1xSPS probes in a largely additive manner. In contrast, the NCM co-activator complex (arrows at right) binds the 1xSPS but not the 2xCSL probe cooperatively. Su(H) concentration increases from 2.5 to 160 nM in 4-fold steps and 2 µM Hairless/NICD/MAM was used in indicated lanes. Note, we separated the two colors for the NCM activating complex in grayscale in D’ and show the entire gel in grayscale in Figure S1A-B. **E**. Average number of sites filled with increasing amounts of Su(H) in reactions of Su(H) alone (black), Su(H) with co-activators (orange) and Su(H) with the Hairless (H) co-repressor (blue). The average number of sites filled is defined as 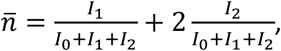 where *I*_0_, *I*_1_ and *I*_2_ are the extracted band values for the 0, 1, and 2 TFs bound to the probe. Each reaction was repeated four times and the dots represent data from each individual experiment. Lines represent fitted data (see methods). Extracted cooperativity factors are as indicated.

To determine the specificity of the engineered synthetic 2xCSL and 1xSPS DNA sequences for Su(H) binding (note that 2xCSL and 1xSPS have the same number of Su(H) binding sites), we performed two tests using electrophoretic mobility shift assays (EMSAs): First, we found that purified Su(H) protein binds DNA probes containing the synthetic 2xCSL and 1xSPS sequences, but not probes with point mutations in the Su(H) binding sites (**Fig 1A-C**). Second, we tested how the orientation of binding sites affects the DNA binding affinity of Su(H) in the presence of NICD and Mam (i.e. the NCM activating complex) or the Hairless (H) co-repressor. For this experiment, we used purified proteins that include the NICD (aa 1763-2412), Mam (aa 87-307) and Hairless (aa 232-358) domains required to form stable complexes with Su(H), and we directly compared the binding of each TF complex using differentially labeled 2xCSL (700nm wavelength, pseudo-colored magenta) and 1xSPS (800nm wavelength, pseudo-colored green) probes in the same reaction (**Fig 1D**). Importantly, we found that like Su(H) alone, the Su(H)/H complex bound both the 2xCSL and 1xSPS probe in an additive manner (**Fig 1D** and **Fig S1A-B**). In sharp contrast, the NCM complex preferentially formed larger TF complexes, consistent with filling both sites of the 1xSPS probe, compared to the sequential binding to 2xCSL (**Fig 1D-D’**). Thus, unlike the NCM co-activator complex, the Su(H)/H repression complex does not bind to SPS sites in a cooperative manner.

To obtain a measure of the cooperativity induced by NCM binding to the 1xSPS vs 2xCSL probes, we quantitatively analyzed the band intensities in the EMSA gels and fitted the extracted values to a 2-site equilibrium binding model (**Fig 1E**). The model takes into account cooperative binding by assuming that the dissociation constant associated with the second binding, *K*_*d*2_, is smaller by a cooperativity factor, C, with respect to the dissociation constant associated with the first binding, *K*_*d*1_, such that *K*_*d*2_ = *K*_*d*1_/*C*. A cooperativity factor higher than 1 corresponds to positive cooperative binding. A cooperativity factor close to 1 or smaller than 1 corresponds to non-cooperative binding and negative cooperative binding (i.e. steric hindrance), respectively. Fitting the band intensities from the EMSA experiments allowed the extraction of the cooperativity factor for each complex and each probe **(Fig S1C-H)**. Other than the NCM complex on SPS, all other experimental conditions exhibited cooperativity factors close to 1, indicating a non-cooperative binding process. In contrast, the NCM complex had a cooperativity factor of 16.9±1.2 on the SPS probe, clearly showing a strong cooperative binding (higher than 16-fold). Taken together, these data show that while the independent sites in the 2xCSL probe mediate similar DNA binding patterns regardless of Su(H) complex, the SPS sites favor the formation of the cooperative NCM activation complexes relative to the binding of Su(H) alone or the Su(H)/co-repressor complex.

### Generation of synthetic SPS and CSL reporters to study Notch-mediated gene regulation in Drosophila

Because each endogenous enhancer integrates transcription factor binding sites (TFBSs) for additional factors and is embedded in its own chromatin environment, it has been difficult to systematically determine how the formation of Notch dimer vs Notch monomer complexes impacts transcriptional outputs. To address this problem in *Drosophila*, we generated fly lines containing transgenic reporters with varying numbers of CSL (*2x, 4x, 8x, or 12xCSL-lacZ*) or SPS (*1x, 2x, 4x or 6xSPS-lacZ*) sites (**Fig 2A**). We first qualitatively characterized *12xCSL-lacZ* and *6xSPS-lacZ* reporter lines inserted into a consistent chromosomal environment using *ϕ*C31 mediated recombination (Bischof et al., 2007) and analyzed for β-gal expression in a variety of tissues. As a control, we generated transgenic reporters containing equal numbers of mutated sites that disrupt Su(H) binding (*12xCSLmut-lacZ* and *6xSPSmut-lacZ*) but left all flanking sequences unchanged (see **Fig 1A-C** for mutant sequence and data indicating a lack of Su(H) DNA binding). Expression analysis in *Drosophila* tissues revealed that the wild type CSL and SPS reporters, but not the mutant reporters, activate qualitatively similar β-gal expression patterns in many Notch-dependent cell types (**Fig 2** and **Fig S2**). For example, both the *12xCSL-lacZ* and *6xSPS-lacZ* transgenes induced reporter expression in the embryonic mesectoderm (**Fig 2B-C**), nervous system, and hindgut dorsal-ventral boundary cells (**Fig 2D-E**). In addition, qualitatively similar expression patterns were observed in the pupal wing disc (**Fig 2H-I**), the pupal eye disc (**Fig 2J-K**), and the adult midgut (**Fig 2L-M**). Surprisingly, however, only the *6xSPS-lacZ* reporter was activated in Notch-active larval imaginal disc cells, such as the wing margin cells (**Fig 2F-G**), the leg joint boundary cells (**Fig S3A-B**), and in differentiating cells of the larval eye (**Fig S3C-D**). In addition, we found that *12xCSL-lacZ* only activated reporter expression in a subset of the cellular patterns generated by the *6xSPS-lacZ* reporter in the larval brain (**Fig S3E-F**).

**Figure 2.**
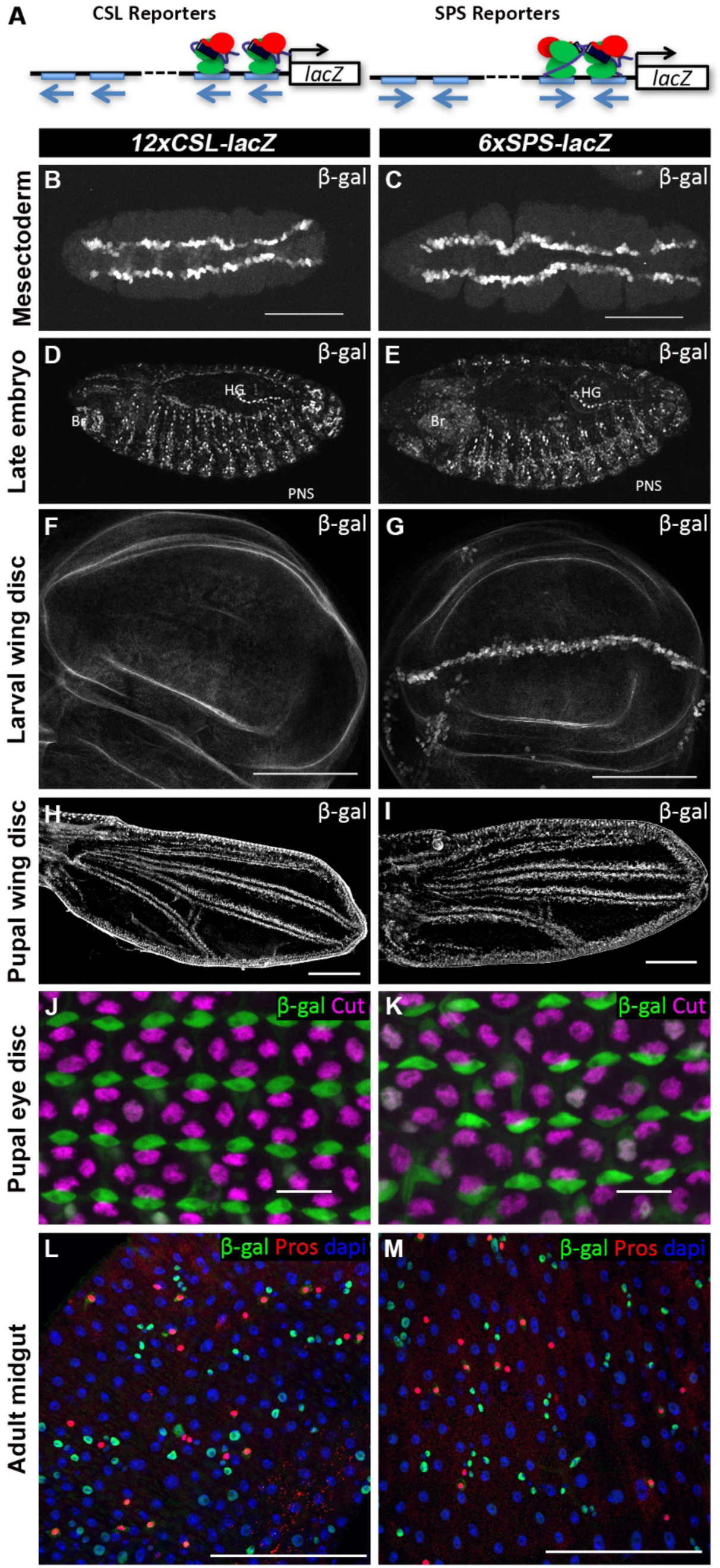
CSL and SPS reporters are expressed in multiple *Notch*-dependent tissues. **A**. Schematics of *NxCSL-lacZ* and *NxSPS-lacZ* reporter constructs. **B-C**. Ventral view of stage 5 *Drosophila* embryos with the *12xCSL-lacZ* (B) and *6xSPS-lacZ* (C) reporters immunostained for β-gal revealed expression in the mesectoderm. **D-E**. Lateral view of stage 15 *Drosophila* embryos with the *12xCSL-lacZ* (D) and *6xSPS-lacZ* (E) reporters immunostained for β-gal revealed expression in expected Notch-dependent tissues including the embryonic brain (Br), peripheral nervous system (PNS), and hindgut (HG). **F-G**. Larval wing discs with the *12xCSL-lacZ* (F) and *6xSPS-lacZ* (G) reporters immunostained for β-gal revealed expression in the D-V boundary cells only with SPS reporter. **H-I**. Pupal wing discs with the *12xCSL-lacZ* (F) and *6xSPS-lacZ* (G) reporters immunostained for β-gal revealed expression in the intervein tissue adjacent to the developing wing veins. **J-K**. Pupal eye discs with the *12xCSL-lacZ* (H) and *6xSPS-lacZ* (I) reporters immunostained for β-gal (green) and cut (magenta), which marks the cone cells, revealed an expression pattern consistent with reporter activity in the primary pigment cells. **L-M**. Adult intestinal midgut cells with the *12xCSL-lacZ* (J) and *6xSPS-lacZ* (K) reporters immunostained for β-gal (green), Pros (red), which is a marker of enteroendocrine cells, and counterstained with DAPI revealed an expression pattern in the smaller nuclei of the midgut, consistent with high Notch activity in the EB cells. Scale bars are 10 μm in J and K, and 100 μm in others.

To investigate if the differential responsiveness of the *6xSPS-lacZ* and *12xCSL-lacZ* transgenes to Notch signaling in larval tissues reflected a position effect, we tested transgenes inserted into an independent chromosomal landing site and found that each was similarly active in the embryo, whereas only *6xSPS-lacZ* was active in the larval wing imaginal disc (**Fig S4**). These data indicate that the failure of the synthetic CSL enhancer to activate reporter expression in larval imaginal disc cells is unlikely due to positional effects of the transgenes. Second, we assessed the activity of the *6xSPS-lacZ* and *12xCSL-lacZ* reporters in embryonic and larval imaginal disc cells that express ectopic NICD. For the *Drosophila* embryo, we used *paired-Gal4* (*prdG4*) to activate a *UAS-NICD* transgene in every other parasegment and found that both *12xCSL-lacZ* and *6xSPS-lacZ* were strongly activated by ectopic NICD (**Fig 3A-B**). In sharp contrast, ectopic NICD expression using *Dpp-Gal4*, which is active in larval wing cells along the anterior-posterior compartment boundary, had a very different impact on these two reporters (**Fig 3C-D**). The *6xSPS-lacZ* reporter was strongly activated by NICD, whereas the *12xCSL-lacZ* reporter showed minimal expression (**Fig 3C-D**). Moreover, it is important to note that the ectopic NICD in this experiment induced ectopic expression of the *cut* Notch target gene and induced pronounced wing overgrowth phenotypes, which strongly suggests that the failure to activate the *12xCSL-lacZ* reporter is not due to limited NICD levels. Third, we used *DppG4* to express a constitutively active Su(H)-VP16 fusion protein, which has been shown to activate Notch target enhancers in an NICD-independent manner (Klein et al., 2000), and found that Su(H)-VP16 activated the *12xCSL-lacZ* reporter in the larval wing disc, although to a lesser extent than it activates *6xSPS-lacZ* (**Fig 3E-F**). Altogether, these data reveal that while synthetic reporters with monomeric CSL sites are sufficient to mediate Notch-dependent expression in embryos, pupal tissues, and the adult midgut, the CSL sites are insufficient to activate strong Notch-dependent processes in larval imaginal disc tissues. While it remains unclear why the CSL reporters fail to respond to NICD in larval imaginal disc cells, the *12xCSL-lacZ* and *6xSPS-lacZ* reporters do elicit qualitatively similar Notch transcriptional responses in many tissues, and thereby provide useful tools to perform quantitative expression analysis between SPS versus CSL reporters inserted into consistent chromosomal locations.

**Figure 3.**
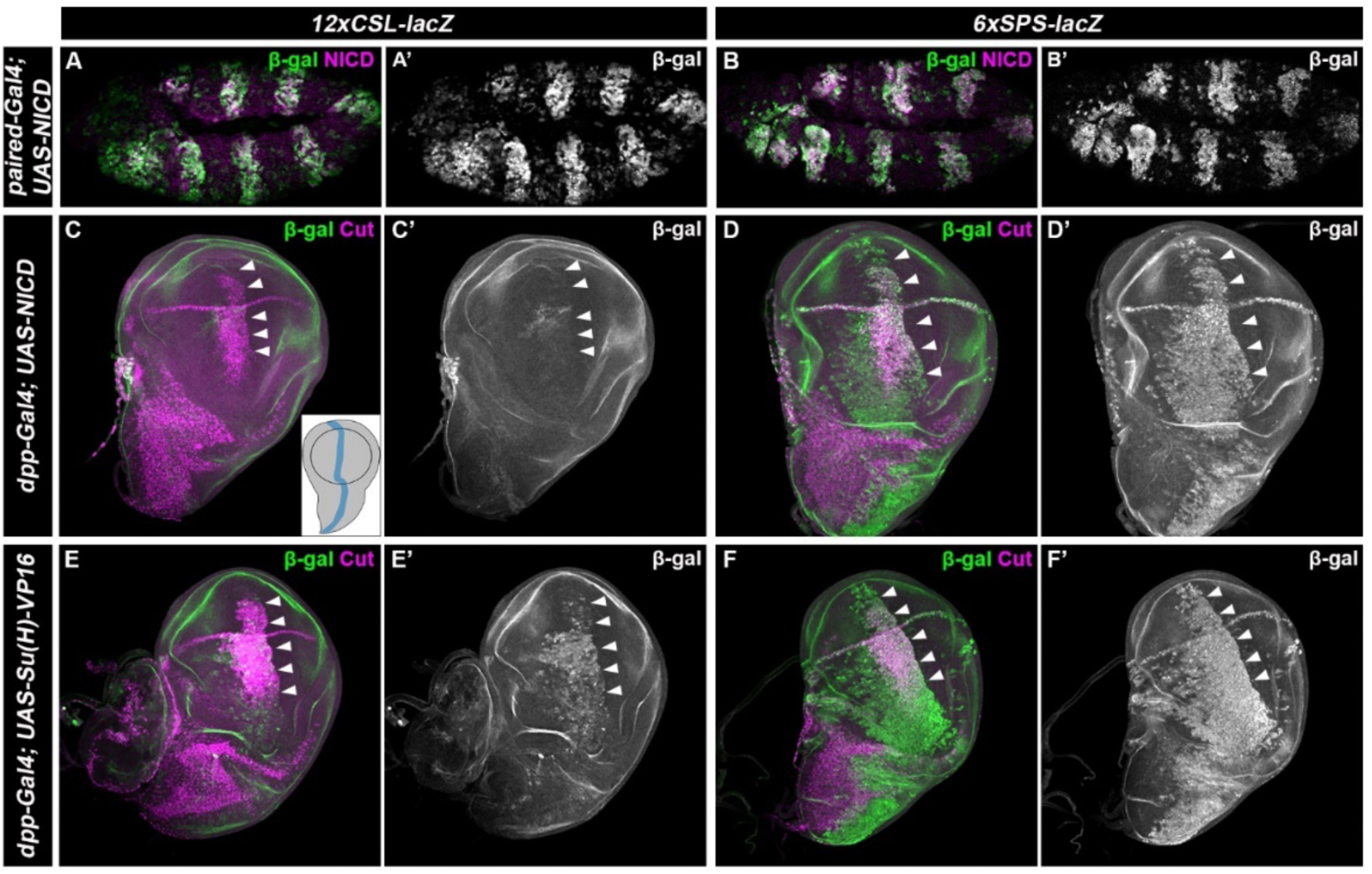
CSL and SPS reporters respond to ectopic Notch signal activation. **A-B**. Stage 11 *Drosophila* embryos containing either the *12xCSL-lacZ* or *6xSPS-lacZ* reporter and *paired-Gal4>UAS-NICD* activation were immunostained for β-gal (green, black and white in A’ and B’) and NICD (magenta). Note, both the *12xCSL-lacZ* and *6xSPS-lacZ* reporters were strongly activated by ectopic NICD. **C-F**. Larval wing discs containing either the *12xCSL-lacZ* or *6xSPS-lacZ* reporter and *dpp-Gal4>UAS-NICD* (C-D) or *dpp-Gal4>UAS-Su(H)-VP16* (E-F) were immunostained for β-gal (green, black and white in C’ and F’) and NICD (magenta). Inset in panel C shows the *dpp-Gal4* positive region in wing discs. Arrowheads denote the region of NICD overexpression in the wing pouch.

### SPS-lacZ reporters exhibit more consistent and stronger response than CSL-lacZ reporters in the mesectoderm

Notch signaling is required for the specification of mesectoderm cell fate by ensuring accurate expression of *single-minded* (*sim*). To assess the ability of both cooperative SPS and independent CSL sites to activate gene expression in the mesectoderm, we analyzed the activity of the *NxCSL-lacZ* and *NxSPS-lacZ* transgenes in the mesectoderm of age-matched embryos using immunofluorescent imaging for both Sim and β-gal protein levels (see methods). Qualitative analysis of reporters containing the same total number of Su(H) binding sites (1xSPS = 2xCSL) revealed that neither a single SPS site (1xSPS-lacZ) nor two CSL sites (2xCSL-lacZ) activated detectable reporter expression in the mesectoderm (**Fig S5**). By contrast, Notch reporter activity in the mesectoderm was observed in embryos containing *lacZ* reporters with 4 or more CSL sites and 2 or more SPS sites (**Fig 4A-F**). These data show that the cooperative binding between NCM complexes does not confer synthetic SPS enhancers with a significantly different response threshold to Notch activation in the mesectoderm from synthetic enhancers with the same number of independent CSL sites.

**Figure 4.**
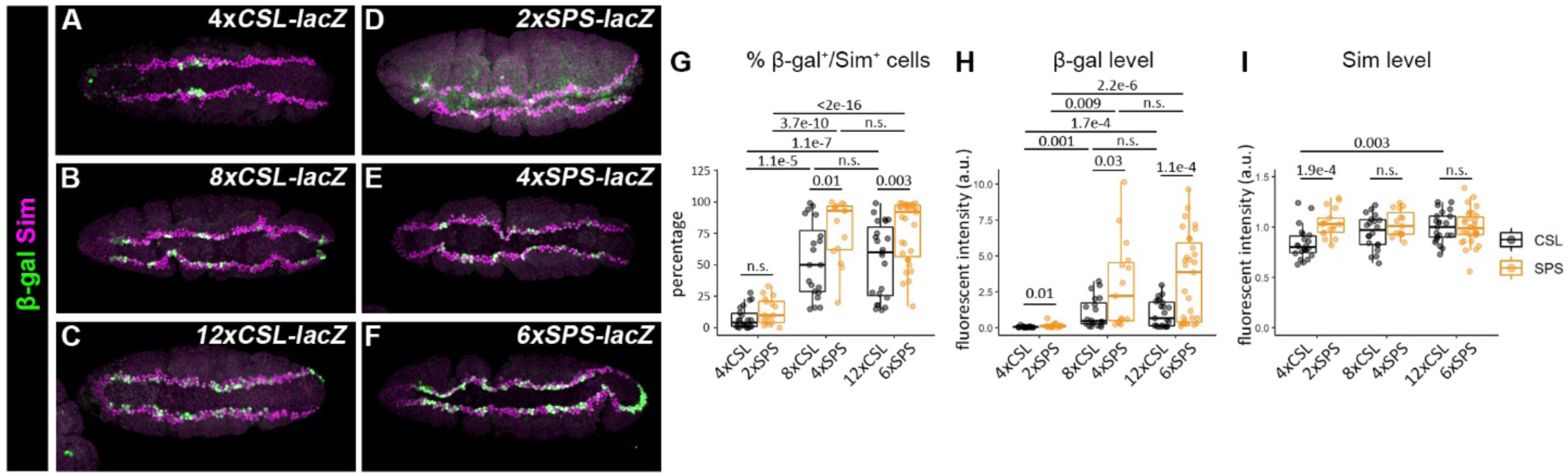
Cooperative binding sites enhance Notch transcriptional activity in the *Drosophila* mesectoderm. **A-F**. Ventral views of stage 5 *Drosophila* embryos carrying either the *4xCSL-lacZ* (A), *8xCSL-lacZ* (B), *12xCSL-lacZ* (C), *2xSPS-lacZ* (D), *4xSPS-lacZ*, (E) or *6xSPS-lacZ* (F) immunostained for β-gal (green) and sim (magenta), which is a marker of mesectoderm cells. **G-I**. Quantification of the percentage of mesectoderm cells (sim-positive cells) that activate β-gal (G), the mean β-gal protein levels (H) and the mean sim protein levels (I) in flies containing the indicated reporters. Each dot represents the measurements from an individual embryo. Box plots show the median, interquartile range, and 1.5 times interquartile range. One-way ANOVA with post-hoc Tukey HSD for equal variance or post-hoc Dunnett’s T3 for unequal variance were used to test significance. n.s. not significant.

Next, we quantitatively assessed how the number and type of binding sites impact transcriptional output in the mesectoderm by analyzing *lacZ* reporter activity in two ways. First, we determined the percentage of mesectoderm cells (as defined by Sim positive staining) that expressed significant levels of β-gal relative to the background (defined as more than 3 standard deviations above the average background fluorescence, see methods) (**Fig 4G**). Second, we measured the intensities of β-gal and Sim in each embryo as a function of binding site type (SPS vs CSL) and binding site number (**Fig 4H-I**). As expected, Sim protein levels did not vary greatly between samples, although a small, but significant difference between the *2xSPS-lacZ* and *4xCSL-lacZ* samples was observed (**Fig 4I**). In contrast, comparative analysis of β-gal expression between these samples revealed the following: 1) Synthetic Notch reporters with SPS sites have a significantly higher likelihood of activating gene expression in each mesectoderm Sim positive cell than synthetic reporters with an equal number of independent CSL sites (**Fig 4G**). For example, the *4xSPS-lacZ* reporter was activated in 78.0±6.3% (mean±sem) of mesectoderm cells in a typical embryo, whereas the *8xCSL-lacZ* reporter was only activated in 53.6±6.7% of mesectoderm cells. Moreover, a similar significant difference was also observed between the *6xSPS-lacZ* and *12xCSL-lacZ* embryos. 2) When comparing synthetic reporters with the same number of Su(H) binding sites (i.e. 8xCSL to 4xSPS), the β-gal levels were significantly higher in the SPS reporter lines than those in the CSL reporter lines (**Fig 4H**). 3) There was a dramatic increase in both the percentage of β-gal-positive/Sim-positive cells (**Fig 4G**) and the levels of β-gal expression (**Fig 4H**) as the number of synthetic binding sites increased from 4xCSL to 8xCSL or from 2xSPS to 4xSPS. However, both the levels of β-gal and the percentage of mesectoderm cells that activated Notch reporter activity were not significantly different between embryos with the *8xCSL-lacZ* and *12xCSL-lacZ* reporters or the *4xSPS-lacZ* and *6xSPS-lacZ* reporters (**Fig 4G-H**), suggesting that Notch-mediated transcriptional activation plateaus above 8 CSL sites and 4 SPS sites. In sum, this analysis revealed that the synthetic SPS reporters are both more likely to be activated and express at higher levels than the synthetic CSL reporters with the same number of binding sites within the Notch-active mesectoderm cells.

### Activation of the SPS reporter gene is more resistant to increased levels of the Hairless co-repressor

Because neither the NICD/Mam co-activators nor the H co-repressor has a DNA binding domain and they bind to Su(H) in a mutually exclusive manner, the activating complexes and repressing complexes likely compete for binding to enhancers to regulate gene expression (Kovall and Blacklow, 2010; Yuan et al., 2016). To determine if the cooperativity between the NCM complex on the SPS results in altered sensitivity to the Hairless co-repressor, we overexpressed Hairless in every other parasegment of stage 11 embryos with *paired-Gal4;UAS-Hairless* and analyzed age-matched embryos for either *12xCSL-lacZ* or *6xSPS-lacZ* reporter activity (see schematic in **Fig 5A**). Interestingly, while similar levels of Hairless overexpression were observed in both reporter lines compared to neighboring non-overexpressing parasegments (**Fig 5B-D**, 2.41±0.08 fold with *12xCSL-lacZ* and 2.43±0.13 fold with *6xSPS-lacZ*, mean±sem), the *12xCSL-lacZ* reporter was more effectively repressed by Hairless overexpression than the *6xSPS-lacZ* reporter (**Fig 5B-C, E**, a 57.5±3.4% reduction in *12xCSL-lacZ* activity versus a 35.0±3.7% reduction in *6xSPS-lacZ* activity compared to the neighboring wild type parasegments, mean±sem). These data are consistent with Su(H)/H complexes more effectively competing with the NCM co-activator for independent CSL sites than for cooperative SPS sites.

**Figure 5.**
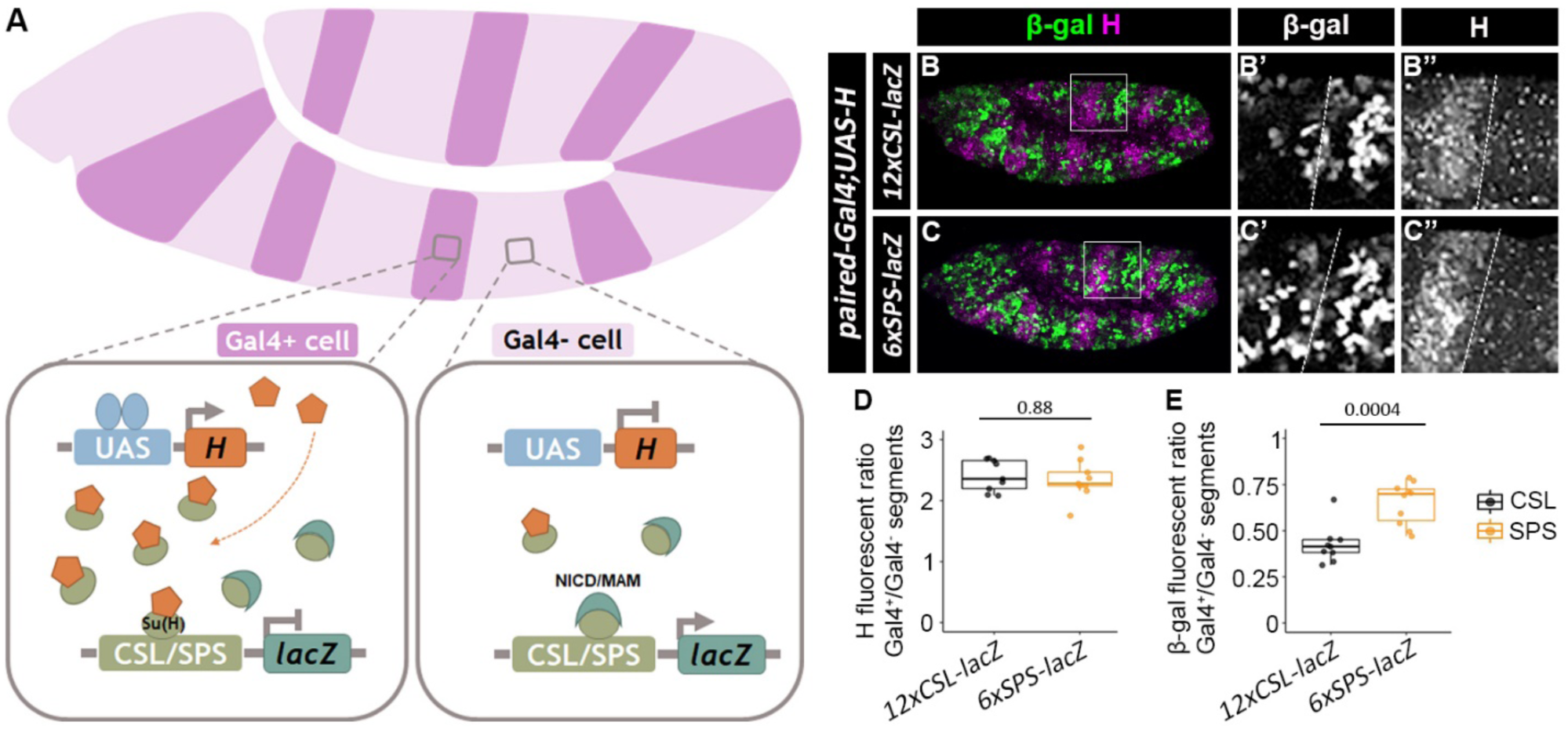
Notch reporters with cooperative binding sites show higher resistance to the Hairless co-repressor than reporters with monomer sites. **A**. Schematic of the over-expression of the Hairless protein using the paired-Gal4>UAS system. Note, that paired-Gal4 is active in every-other parasegment and thereby allows the direct comparison of Gal4-positive (Gal4+) regions that express endogenous and exogenous Hairless with wild type (Gal4-) regions that only express endogenous Hairless in the same embryo. **B-C**. Lateral views of stage 11 *paired-Gal4>UAS-Hairless* embryos containing either the *12xCSL-lacZ* (B) or *6xSPS-lacZ* (C) reporter. Embryos were immunostained with β-gal (green) and Hairless (magenta) and close-up views of the individual channels in black and white for the highlighted regions are shown in B’-C’ (β-gal) and B’’-C’’ (Hairless). **D-E**. Quantification of ratios of Hairless (D) and β-gal (E) in parasegments with ectopic Hairless (*paired-Gal4*^*+*^) compared to control parasegments (*paired-Gal4*^*-*^). Each dot represents the mean measurement from an individual embryo containing either the *12xCSL-lacZ* or *6xSPS-lacZ* reporter. Box plots show the median, interquartile range, and 1.5 times interquartile range. One-way ANOVA was used to test significance.

An alternative explanation for the increased resistance to Hairless of the *6xSPS-lacZ* reporter is that when both Su(H)/H and NCM activation complexes are bound to neighboring sites on the same enhancer, the Su(H)/H complex may more efficiently antagonize the activation potential of the monomer NCM activation complex than that of the dimer NCM complex. To test this idea, we targeted the Hairless co-repressor to heterologous DNA binding sites using 5 copies of the LexA DNA binding site (5xLexAop) inserted adjacent to either 12xCSL or 6xSPS binding sites (**Fig 6A**). To do so, we generated a *UAS-V5-LexADBD-Hairless*^*Δ232-263*^ construct and overexpressed this fusion protein with *paired-Gal4* in reporter lines containing LexA operator binding sites (i.e. *5xlexAop-12xCSL-lacZ* or *5xlexAop-6xSPS-lacZ*). Deletion of the Hairless Δ232-263 amino acids removes the Su(H)-binding domain (Maier et al., 2011), and thus renders this protein incapable of being recruited to CSL or SPS sites. Hence, overexpressing the V5-LexADBD-H^Δ232-263^ protein had negligible impacts on the expression of the *12xCSL-lacZ* and the *6xSPS-lacZ* reporters lacking lexAop sites (**Fig 6B-C, F**). In contrast, expressing the V5-LexADBD-H^Δ232-263^ protein strongly repressed the activity of both the *5xlexAop-12xCSL-lacZ* and the *5xlexAop-6xSPS-lacZ* reporters to a similar degree (**Fig 6D-F**). As additional controls, expressing a V5-lexADBD protein that lacks the Hairless protein or expressing the V5-Hairless^Δ232-263^ protein that is not targeted to DNA failed to repress the *5xlexAop-12xCSL-lacZ* and *5xlexAop-6xSPS-lacZ* reporters (**Fig S6**). Altogether, these data suggest that when Hairless is specifically targeted to DNA sites near where the NCM complex binds, it efficiently antagonizes NCM mediated activation regardless of site architecture. However, in wild type embryos where the Hairless/Su(H) complex and the NCM complex compete for binding sites, the cooperativity of the NCM complex for SPS sites makes these synthetic enhancers more resistant to Su(H)/H binding and repression.

**Figure 6.**
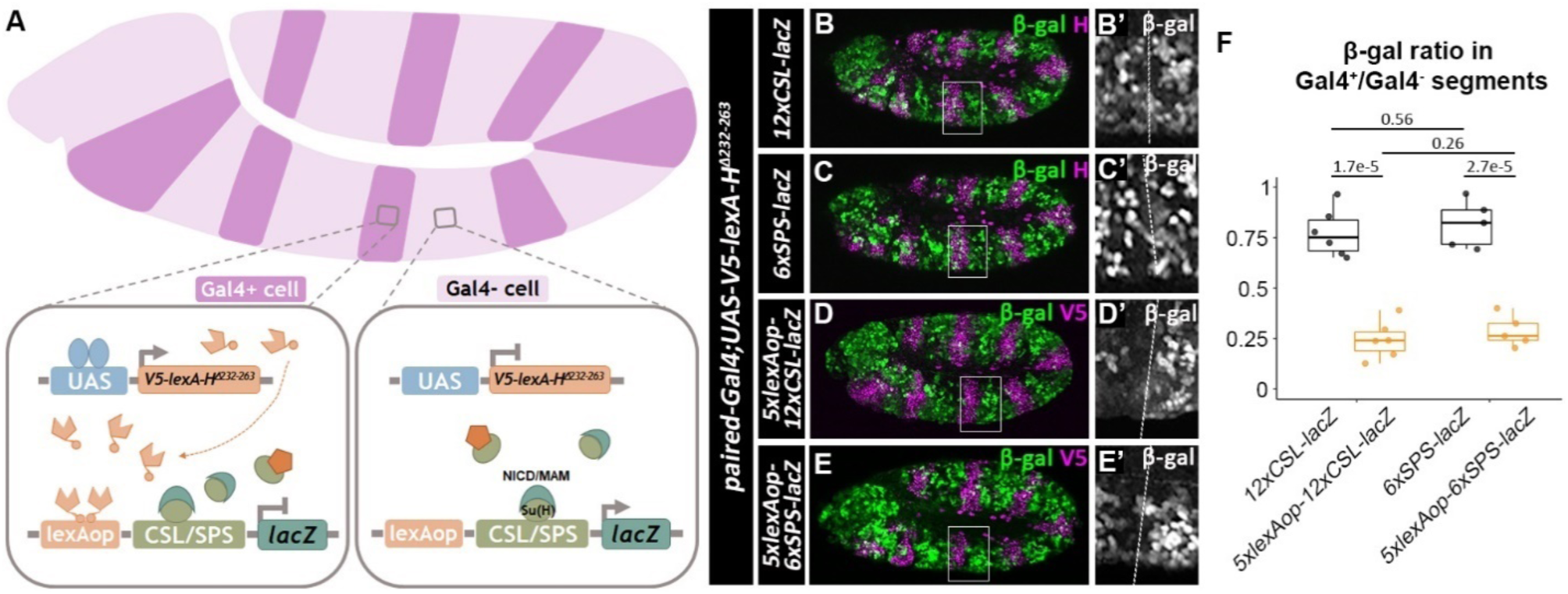
Hairless represses both cooperative and non-cooperative Notch-mediated transcriptional activation when targeted to DNA via a heterologous DNA binding domain. **A**. Schematic of the over-expression of a V5-lexA-HairlessΔ232-263 protein using the paired-Gal4-UAS system. Note, that the Hairless Δ232-263 deletion removes the protein domain that interacts with Su(H). Thus, this protein neither directly competes with NICD/Mam for binding to Su(H) nor does it get recruited to the CSL/SPS binding sites. Instead, the V5-LexA-HΔ232-263 protein is targeted to DNA via lexAop sequences that have been inserted into the CSL/SPS reporter vectors. **B-C**. Stage 11 embryos of *paired-Gal4>UAS-V5-lexA*^*DBD*^*-Hairless*^*Δ232-263*^ immunostained with β-gal (green) and Hairless (magenta) with either *12xCSL-lacZ* or *6xSPS-lacZ* reporter. A’-B’. Close-up views of β-gal intensity in black and white are shown in insets from A-B with the *paired-Gal4*-positive parasegment on the left and the *paired-Gal4*-negative parasegment on the right. **D-E**. Stage 11 embryos of *paired-Gal4>UAS-V5-lexA*^*DBD*^*-Hairless*^*Δ232-26*^ immunostained with β-gal (green) and V5 (magenta) with either *5xlexAop-12xCSL-lacZ* or *5xlexAop-6xSPS-lacZ*. C’-D’. Close-up views of β-gal intensity in black and white are shown in insets from C-D with the *paired-Gal4*-positive parasegment on the left and the *paired-Gal4*-negative parasegment on the right. **F**. Quantification of ratios of β-gal of paired-Gal4^+^ to paired-Gal4^-^ parasegments in flies with indicated genotypes. Each dot represents the average measurement from an individual embryo containing the indicated reporter. Box plots show the median, interquartile range, and 1.5 times interquartile range. One-way ANOVA used to test significance.

## Discussion

In this study, we investigated how differences in DNA binding site architecture (CSL vs SPS) impact the DNA binding of the *Drosophila* Su(H) co-activator and co-repressor complexes *in vitro* and transcriptional output *in vivo*. Using a combination of *in vitro* DNA binding assays, synthetic biology, and *Drosophila* genetics, we made three key findings that reveal new insights into the differences between monomeric CSL sites and dimeric SPS sites in mediating Notch-dependent transcription. First, we found that unlike the Su(H)/NICD/Mam activating complex, the tested Su(H)/H repressor complex does not interact with SPSs in a cooperative manner and instead binds in a similar additive manner to both CSL and SPS probes. Second, we found that while transgenic reporter genes containing the same number of CSL vs SPS binding sites largely mediate the same qualitative expression patterns in *Drosophila* (with the noted exception of larval tissues), the synthetic SPS enhancers are more consistently activated and activated to a higher level in the mesectoderm relative to the synthetic enhancer with equal numbers of monomeric CSL sites. Third, we found that the Hairless co-repressor can more readily repress Notch induced activation of the synthetic CSL enhancers than the synthetic SPS enhancers, and this effect was only seen when both NICD and H bind the enhancer via Su(H). Overall, these data support the model that, compared to enhancers with only CSL sites, Notch-regulated enhancers with cooperative SPSs will more likely be bound by the co-activator complex and thereby are more resistant to the potential negative impacts of CSL/co-repressor complexes. Below, we integrate these findings with other publications on Su(H) stability, Notch transcriptional dynamics, and endogenous Notch-regulated enhancers.

Recent studies in *Drosophila* have demonstrated that the Su(H) TF is unstable in the absence of either the Notch signal (NICD) or the Hairless co-repressor (Praxenthaler et al., 2017). Moreover, biochemical assays demonstrated that Su(H), as well as the mammalian RBPJ CSL TF, uses distinct but overlapping domains to bind NICD and co-repressors and do so with similar affinities (Collins et al., 2014; Friedmann et al., 2008; Yuan et al., 2016, 2019). Together, these findings suggest that the NICD/Mam co-activator proteins and the Hairless co-repressor protein compete to bind Su(H) in a mutually exclusive manner, and that the vast majority of Su(H) in a cell is in either an activating or repressing complex. Hence, Notch-mediated transcriptional output is dependent upon which TF complex interacts with the binding sites found in Notch-regulated enhancers. Our DNA binding data show that monomeric CSL sites bind similarly to both the Su(H)/NICD/Mam activating complex and the Su(H)/H repressing complex. In contrast, the paired sites found in SPS enhancers cooperatively bind the Su(H)/NICD/Mam complex, but not the Su(H)/H co-repressor complex. In fact, our biochemical studies show that the effective Kd for binding a second co-activator complex to the SPS site is ∼17 times smaller than the effective Kd of binding a second Su(H)/H co-repressor complex or a second Su(H) TF alone to the SPS site. However, since a truncated Hairless protein was used in our EMSAs, we cannot rule out the possibility that regions outside of the tested construct contribute to cooperativity. But importantly, the DNA binding data are congruent with the stronger and more consistent reporter expression driven by the SPS enhancers in the mesectoderm as compared to CSL reporters integrated into the same chromosomal locus.

Previous studies on Notch-dependent transcription in the mesectoderm used the MS2-MCP-GFP system to characterize the transcriptional dynamics of two enhancers: the *E(spl)m5/m8* mesectoderm enhancer (MSE) that has an SPS site as well as several potential monomeric CSL sites, and the *sim* MSE enhancer that lacks SPSs but contains monomeric CSL sites (Falo-Sanjuan et al., 2018). Interestingly, the *E(spl)m5/m8* MSE and *sim* MSE enhancers showed very similar transcriptional dynamics and highly correlated transcriptional activity, suggesting that SPS and CSL sites mediate similar transcriptional responses within the mesectoderm. However, when different doses of ectopic NICD were provided in neighboring cells, the *E(spl)m5/m8* MSE enhancer drove expression significantly earlier than the *sim* MSE, consistent with the notion that the *E(spl)m5/m8* MSE enhancer displays a lower detection threshold for NICD to activate transcription. Intriguingly, this difference in enhancer activity was likely due to additional TF inputs and not due to the SPS site, as neither converting it into 2 CSL sites within the *E(spl)m5/m8* MSE nor adding an SPS site to the *sim* MSE changed the timing of their activity. In contrast, both the SPS-containing wild type *E(spl)m5/m8* MSE enhancer and the engineered SPS-containing *sim* MSE enhancer were found to activate higher levels of gene expression due to increased transcriptional burst size.

Our findings using synthetic enhancers, which unlike the more complex endogenous enhancers use isolated SPS and CSL sites, are largely in agreement with the results obtained using live imaging in *Drosophila* embryos. First, we found that synthetic SPS and CSL enhancers both required the same number of Su(H) binding sites (*2xSPS* vs. *4xCSL*) to activate reporter expression within the mesectoderm, whereas the *1xSPS-lacZ* and *2xCSL-lacZ* reporters both failed to activate gene expression in the mesectoderm. This finding is in line with SPS and CSL sites having similar NICD detection thresholds. Second, we found that SPS enhancers activated transcription more consistently and at a higher level than CSL enhancers with the same total number of binding sites. Third, we investigated if the SPS-mediated cooperativity grants the *Drosophila* co-activators any advantages over the co-repressors and found that only the NCM complex, but not the Su(H)/H co-repressor complex, cooperatively binds SPSs. Consistent with the idea that cooperative binding to SPSs may lead to increased resistance to changes in co-repressor levels, our reporter assays showed that when Hairless was overexpressed at a moderate level (∼2 fold overexpression) and had to compete with co-activators for Su(H) binding, the SPS reporter was more resistant to Hairless than the CSL reporter. However, when we targeted Hairless to DNA via an independent non-competitive mechanism, the Hairless co-repressor was equally competent to antagonize the activation effects elicited by the NCM complex on either the synthetic CSL or SPS reporters. Thus, the co-activator and co-repressor complexes compete for binding sites, and the cooperativity of the NCM to SPSs results in a competitive advantage for the activation complex over the repression complex.

Integrating our findings using the synthetic SPS and CSL enhancers with the studies on transcription dynamics of endogenous enhancers supports the following model: The cooperative binding of NCM activating complexes on SPSs results in enhanced stability of the NCM complex (i.e. a slower off-rate) relative to independent CSL sites, which induces larger transcriptional burst sizes and enhanced levels of gene expression. In addition, cooperative NCM binding to SPS enhancers renders these binding sites less sensitive to the repressive impacts of the Su(H)/Hairless co-repressor complex. Importantly, each of these properties (i.e. cooperative NCM binding and preferential co-activator binding to DNA over co-repressor binding) would result in more consistent and higher transcription levels of target genes. However, there are several factors to consider regarding how these differences in synthetic SPS versus CSL enhancer activity can be translated to endogenous Notch regulated enhancers. First, most endogenous enhancers with SPS also contain one or more independent CSL sites that are more highly sensitive to the Hairless co-repressor. Hence, the transcriptional dynamics and ultimate output of endogenous Notch enhancers are likely to be influenced by the combined number and accessibility of the SPS versus CSL binding sites. Second, we previously found that synthetic SPS enhancers, but not CSL enhancers, can induce a *Notch* haploinsufficiency phenotype via a CDK8-dependent NICD degradation mechanism (Kuang et al., 2020). This finding suggests that SPSs preferentially promote NICD turnover, and such activity has potential implications for Notch-dependent transcriptional dynamics. Third, while we found that synthetic SPS and CSL reporters largely activate similar expression patterns in a variety of Notch-dependent tissues, only the SPS enhancers mediated significant Notch-dependent output in larval imaginal tissues. Intriguingly, this unexpected finding is consistent with a published study showing that an SPS-containing *E(spl)m8* enhancer that activates Notch-dependent gene expression in dorsal-ventral boundary cells of the larval wing disc fails to activate reporter gene expression when the orientations of the Su(H) sites were reverted from an SPS to non-cooperative CSL sites (Cave et al., 2005). These findings suggest that CSL sites alone are insufficient to activate Notch-dependent outputs in larval tissues without the influence of additional TF inputs. Thus, additional studies focused on using enhancers with distinct compositions of binding sites in different Notch-dependent tissues are needed to further understand how the NCM activation and CSL/co-repressor complexes are integrated with additional transcriptional inputs to mediate cell-specific output.

## Supporting information

Supplemental Figures-Kuang et al

Supplemental Tables-Kuang et al

## Acknowledgements

We thank members of the Gebelein lab for comments on this work. We thank Stephen Crews (University of North Carolina) for the kind gift of the *sim* cDNA, and Anette Preiss and Dieter Maier (University of Hohenheim) for the kind gift of the Hairless antibody. We also thank Thomas Klein for the kind gift of the UAS-Su(H) fly line (University of Dusseldorf) and the *Drosophila* stock centers and Developmental Studies Hybridoma Bank (DSHB) for fly stocks and antibody reagents.

## Contributions

This scientific study was conceived and planned by Y.K., R.K., D.S., and B.G. The synthetic enhancers were designed by M.T.W. and B.G. The transgenic constructs and *Drosophila* reagents were generated by Y.K., B.C., A.P., and B.G. The *Drosophila* experiments were performed by Y.K., L.M.G., and A.P. Protein purification and gel shift analysis was performed by Y.K., E.K., and R.A.K. Data analysis was performed by Y.K., N.E., O.A., R.K., D.S., and B.G. The mathematical modeling was performed by N.E., O.A., and D.S. The manuscript was written by Y.K. and B.G.

## Materials and Methods

### Protein purification and electrophoretic mobility shift assays (EMSAs)

*Drosophila* proteins used in EMSAs include Su(H) (aa 98-523), Hairless (aa 232-358), NICD (aa 1763-2412) and Mastermind (aa 87-307). Recombinant proteins of each were expressed in *E. coli* and purified using affinity (Ni-NTA or Glutathione) ion exchange and size exclusion chromatography as previously described (Friedmann, et al., 2008). The purity of proteins was determined by SDS-PAGE with Coomassie blue staining and protein concentration was measured by UV280 absorbance. EMSAs were performed as previously described (Uhl, et al., 2016; Uhl, et al., 2010). Fluorescent labeled probes were mixed with purified proteins and incubated at room temperature for 20 minutes before loading. The protein concentration used for each experiment is listed in each Figure legend. Probe sequences are listed in Supplementary Table 1. Acrylamide gels were run at 150V for 2 hours and then imaged using the LICOR Odyssey CLx scanner.

### EMSA quantification

The raw data for the mathematical analysis was extracted from gray scale images of the EMSA gels. The entire process was performed with custom MATLAB code. We utilized a local minima algorithm to extract the inter-lane intensity values. Inter-lane values were used to fit the appropriate background value to a specific location in the image. Band values were extracted by calculating the background subtracted intensity sum over rectangular boxes, which were optimized for maximal signal to background.

We then fitted the extracted band intensities to a model for binding to two sites that takes into account cooperative binding to the second site. The binding probability is calculated using standard equilibrium binding kinetics (Michaelis-Menten) to the two sites. Cooperativity is introduced into the model by assuming that the binding dissociation constant of an activation or repression complex to the second site is reduced by a cooperativity factor *C*, namely that 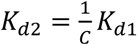. The corresponding probabilities that the probe is bound by 0, 1 or 2 complexes are given by:

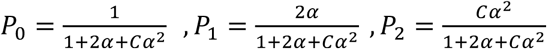

where 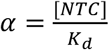 is the statistical weight associated with binding of a complex to a CSL or SPS site. *K*_*d*_ is the equilibrium dissociation constant to a single site. If the value of *C* is equal to 1, then the binding to the two sites is non-cooperative. If *C* > 1 then the cooperativity is positive (2^nd^ binding is enhanced). If *C* < 1 then the cooperativity negative (2^nd^ binding is suppressed).

We observed that even at high concentrations of Su(H) the 1-site state is never depleted (e.g. see NCM on SPS), and the signal of the 0-site state never decays to zero. We therefore assumed that there is a probability-*f* that a site will become unavailable for binding. Under this assumption there is a fraction *f*^2^ of the probes that will have no functioning sites (i.e. that both sites are unavailable), and a fraction 2*f* of the probes that have only 1 functioning site (one of the two sites is unavailable). In this case the probability to find the probe is modified to:

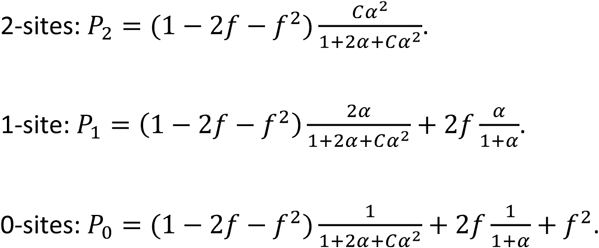

We then fit the normalized band intensities using least mean square to the sum of these three expressions. The fitting parameters are *K*_*d*_, *C*, and *f*. The parameters are extracted for each experiment separately. The confidence interval on the fitting parameters was calculated using a bootstrap method where 5000 random data sets with the same mean and standard deviation as those observed experimentally were generated. The fitting procedure was then applied to all bootstrapped data to obtain the distribution of fitting parameters. The confidence intervals were determined by calculating the 95-percentile range for each parameter.

### Generation of transgenic flies

12xCSL and 6xSPS synthetic enhancers were designed and synthesized as previously reported (Kuang et al., 2020). The 2xCSL, 4xCSL, 8xCSL, 1xSPS, 2xSPS, and 4xSPS sequences were synthesized as oligonucleotides containing appropriate restriction enzyme site overhanging sequences to aid cloning into the *placZ-attB* vector (Bischof et al., 2007). The 5xlexAop sequence was synthesized by Genscript with flanking HindIII and EcoR1 restriction enzyme sites to aide cloning into the following vectors: *placZ-attB*; *12xCSL-lacZ* or *6xSPS-lacZ*. The coding sequences for the LexA-DBD and Hairless-Δ232-263 sequences were synthesized by Genscript with appropriate flanking restriction enzyme sites for cloning into a modified pUAST vector that contained an N-terminal V5-epitope tag. These synthesized DNAs were used to generate the pUAST-V5-lexADBD, pUAST-V5-HairlessΔ232-263, and pUAST-V5-LexADBD-HairlessΔ232-263 vectors. All sequences were confirmed by Sanger sequencing, purified using Qiagen Midi-prep Kit and sent for *Drosophila* injection to Rainbow Transgenic, Inc. Transgenic *Drosophila* lines were established by integration into the *Drosophila* genome using phiC31 recombinase integrase and landing sites located at either 51C or 86Fb as indicated (Bischof et al., 2007). All newly derived sequences and restriction sites used for cloning are listed in Supplementary Table 2.

### Fly husbandry

The following alleles were obtained from the Bloomington *Drosophila* Stock Center: paired-Gal4 (#1947), UAS-NICD (#52008), and UAS-Hairless (#15672) and the UAS-Su(H)-VP16 line was previously described (Kidd et al., 1998). Flies were maintained at 25°C and under standard conditions.

### Generation of single-minded (sim) antibody

Guinea pig anti-Sim serum was generated as previously described (Gutzwiller et al., 2010). Briefly, a Sim cDNA was gifted from Dr. Stephen Crews (University of North Carolina). The cDNA sequence corresponding to sim-PD (aa 361-672) was PCR amplified and cloned in-frame with a 6xHis-Tag into a modified pET-14b plasmid (Novagen). The expression plasmid was transformed into BL21 competent *E. coli* and the expression of the fusion protein was induced by IPTG. The His-tag-Sim protein was extracted in 8M urea lysis buffer, purified by Ni-NTA affinity chromatography, confirmed by Coomassie blue staining, and injected into guinea pigs to generate anti-Sim serum (Cocalico Biologicals, Inc).

### Immunostaining and quantitative analysis

For mesectoderm cells, 2-4 hour-old *Drosophila* embryos of the indicated genotypes were collected, fixed, and immunostained using anti-β-gal (chicken 1:1000, Abcam ab9361) and anti-Sim (guinea pig 1:500, this study) serum and appropriate secondary antibodies conjugated to fluorescent dyes (Jackson Labs). Age-matched embryos were selected and imaged under identical settings using a Nikon A1R inverted confocal microscope (20x objective). The mesectoderm cells were defined as Sim-positive and selected for quantitative expression analysis. To do so, pixel intensity of both Sim and β-gal was subsequently determined with background correction using Imaris software. β-gal positive cells were defined as cells with pixel intensity at least three-fold higher than the standard deviation of the background measurements. One-way ANOVA with proper post-hoc tests was used to determine statistical significance.

For the *paired-Gal4* experiments, 0-16 hour-old embryos were collected, fixed, and immunostained with either Hairless (guinea pig 1:500, Annett and Dieter) and β-gal or V5 (mouse 1:500, Invitrogen R960-25) and β-gal as indicated. Stage 11-12 *Drosophila* embryos were imaged under identical settings in each experiment by either a ZEISS Apotome or Nikon A1R inverted confocal microscope. Fluorescent intensity was quantified using Fiji software as previously described (Zandvakili et al., 2018, 2019). Briefly, the z-stack images were sum-projected and the Gal4^+^ and Gal4^-^ regions in embryos were manually determined. The ratio of β-gal or Hairless was calculated between the Gal4^+^ and Gal4^-^ parasegments after background subtraction. One-way ANOVA with proper post-hoc tests was used to determine statistical significance. The larval wing discs were dissected, fixed, and stained as described (Kuang et al., 2020). Pupal eye discs were fixed for 30 min, and pupal wing discs and adult posterior midguts were fixed for 45 min in 4% formaldehyde after dissection. All of the samples were stained as previously described (Kuang et al., 2020). Antibodies used in this study include β-gal (chicken 1:1000, Abcam ab9361) and sim (guinea pig 1:500, this study), cut (mouse 1:50, DSHB 2B10), Pros (mouse 1:100, DSHB MR1A), Hairless (guinea pig 1:500, Annett and Dieter) and V5 (mouse 1:500, Invitrogen R960-25).

