## Supplemental Figures-Kuang et al for "Enhancers with cooperative Notch binding sites are more resistant to regulation by the Hairless co-repressor"

**Kuang et al Supplemental Figures:**

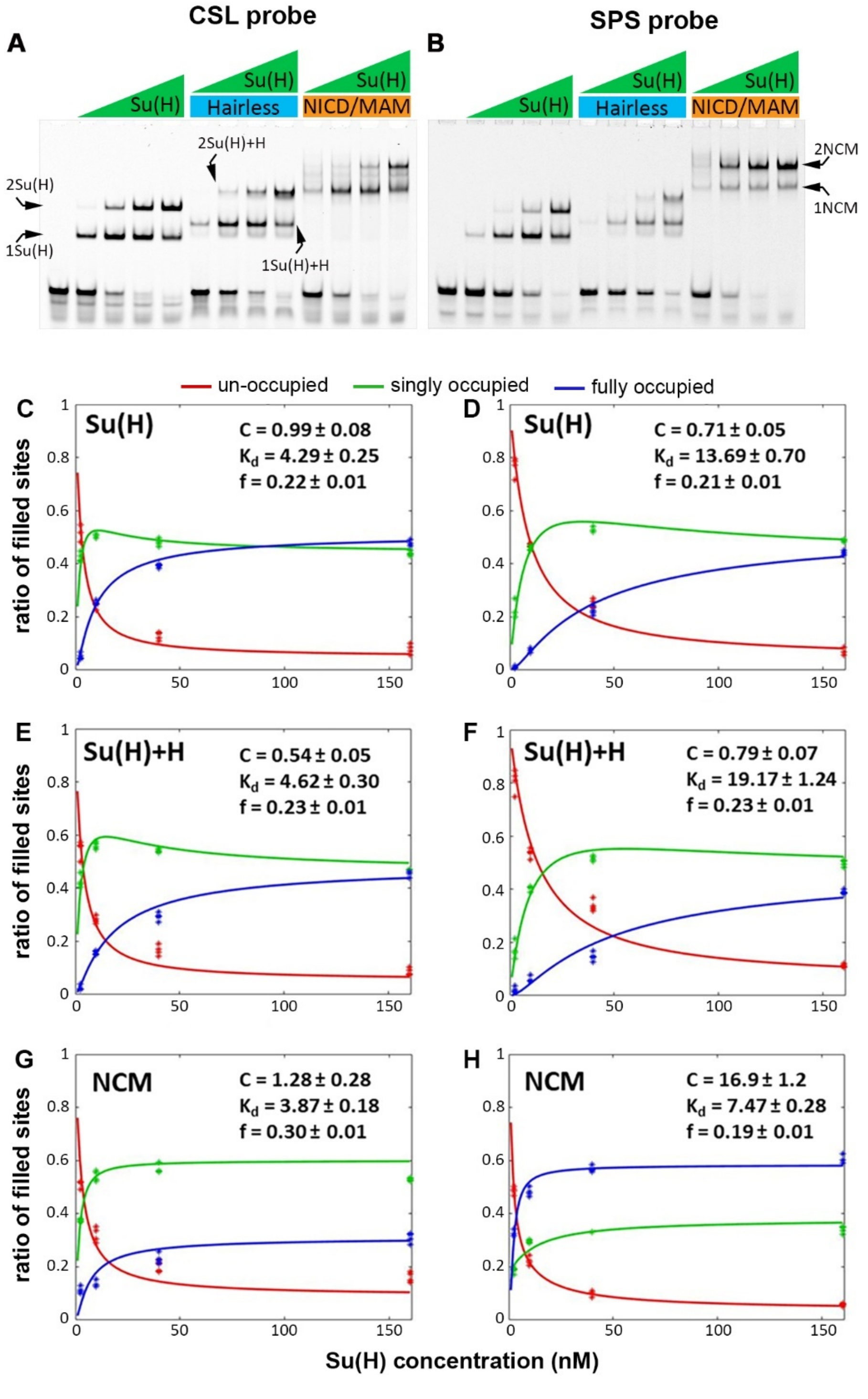


**Figure S1. EMSA quantification and modeling of DNA binding states. A-B**. Individual channels of the same EMSA data shown in Figure 1D. **C-H**. Quantification of the amount of probe that was not bound (unoccupied, red line), bound by a single complex (green line), and bound by two complexes (fully occupied, blue lines). The data for the 2xCSL probe is shown at left, whereas the data for the 1xSPS probe is shown at right. The concentration of Su(H) used is shown along the X-axis. **C-D**, Su(H) was added to each reaction in the absence of either the co-activators or co-repressors. **E-F**, Su(H) was added to each reaction with an excess of the Hairless co-repressor**. G-H**, Su(H) was added to each reaction with an excess of NICD and Mam (NCM). Data points were extracted from EMSAs and represented as asterisks. Simulated data from the model are represented in lines. *C*, cooperativity factor. $K_d$, equilibrium dissociation constant. f, fraction of sites unavailable for binding. Data are from four EMSA gels.

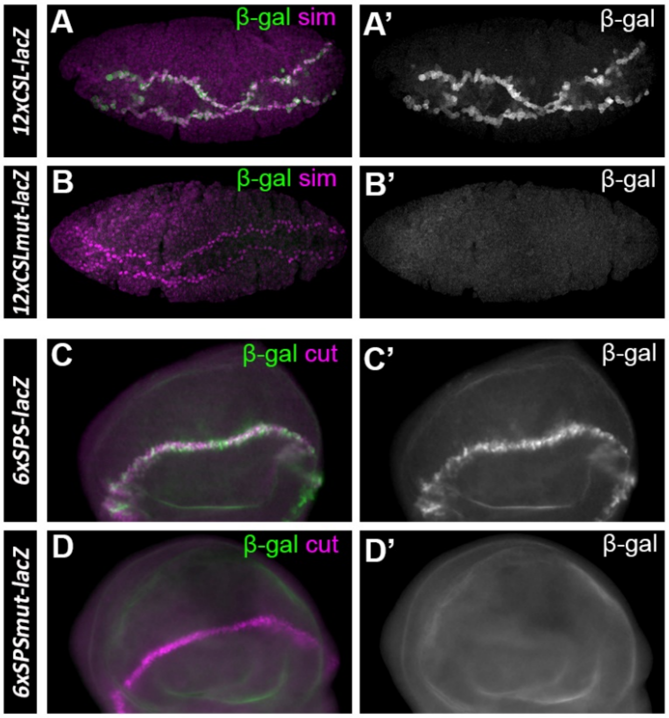


**Figure S2. Mutating the Su(H) binding sites abolishes transcriptional activity from the CSL and SPS reporters in *Drosophila* tissues. A-B**. Stage 5 *Drosophila* embryos containing either the *12xCSL-lacZ* or the *12xCSLmut-lacZ* reporter were immunostained and imaged under identical conditions for β-gal (green, black and white in A' and B') and sim (magenta). Note, the mesectoderm expression activity of the *12xCSL-lacZ* reporter is lost when the CSL binding sites were mutated. **C-D**. Larval wing discs containing either the *6xSPS-lacZ* or the *6xSPSmut-lacZ* reporter were immunostained and imaged under identical conditions for β-gal (green, black and white in C' and D') and cut (magenta). Note, the wing margin cell expression activity of the *6xSPS-lacZ* reporter is lost when each SPS binding site was mutated.

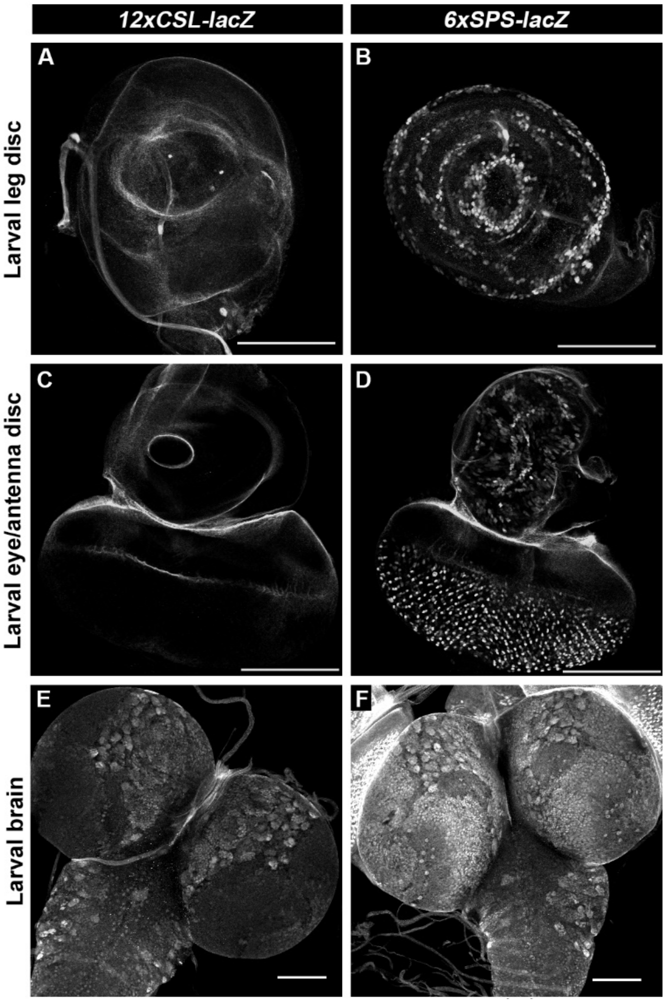


**Figure S3. CSL and SPS reporters significantly differ in expression activity in larval imaginal disc and brain tissues.** β-gal immunostaining of third instar larval imaginal discs reveals that the *6xSPS-lacZ* reporter, but not the *12xCSL-lacZ* reporter, is active in the expected pattern in larval leg discs (**A-B**), larval eye-antenna discs (**C-D**). In contrast, β-gal expression is observed in *12xCSL-lacZ* larval brain tissue (**E**), although in a more limited pattern than in the 6xSPS-lacZ larval brain (**F**). Images were taken under the same setting. Scale bar, 100 µm.

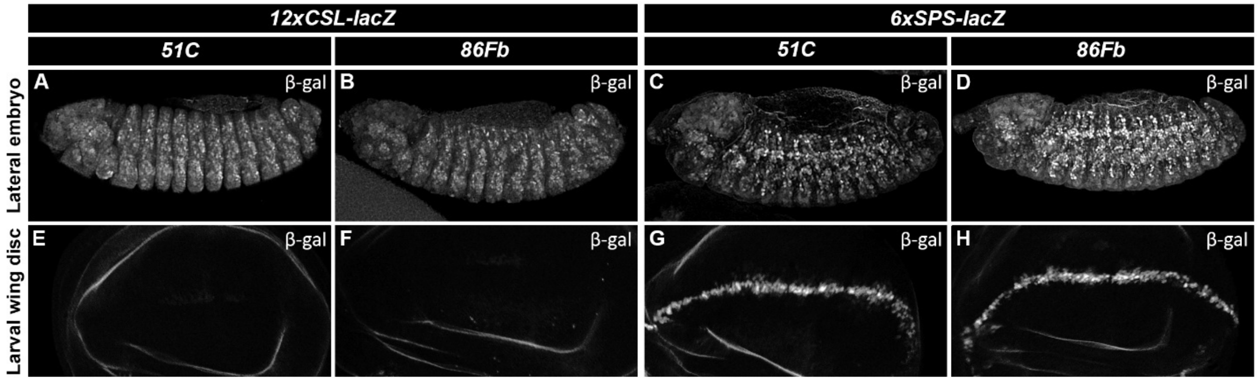


**Figure S4. *12xCSL-lacZ* and *6xSPS-lacZ* transgenic reporters behave similarly in two different loci. A**-**D**. Stage 15 *Drosophila* embryos homozygous for either the *12xCSL-lacZ* (A-B) or *6xSPS-lacZ* (C-D) at the indicated genomic loci (51C or 86Fb) were immunostained with β-gal. Note, the similar expression patterns by both transgenes in each chromosomal location. **E**-**H**. Third instar larval wing imaginal discs homozygous for the indicated reporters were immunostained with β-gal. Note, only the 6xSPS-lacZ reporter is active in the wing margin cells, whereas the *12xCSL-lacZ* fails to activate significant gene expression when inserted into either the 51C or 86Fb locus.

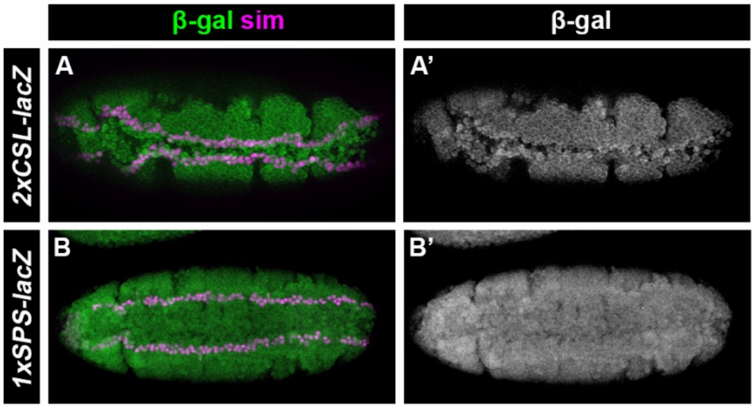


**Figure S5. 2xCSL and 1xSPS reporters fail to mediate Notch activation in mesectoderm cells. A-B**. Stage 5 *Drosophila* embryos containing either the *2xCSL-lacZ* (A) or *1xSPS-lacZ* (B) reporter were immunostained and imaged under identical conditions for β-gal (green, black and white in A' and B') and Sim (magenta). Note, neither reporter activates in the mesectoderm.

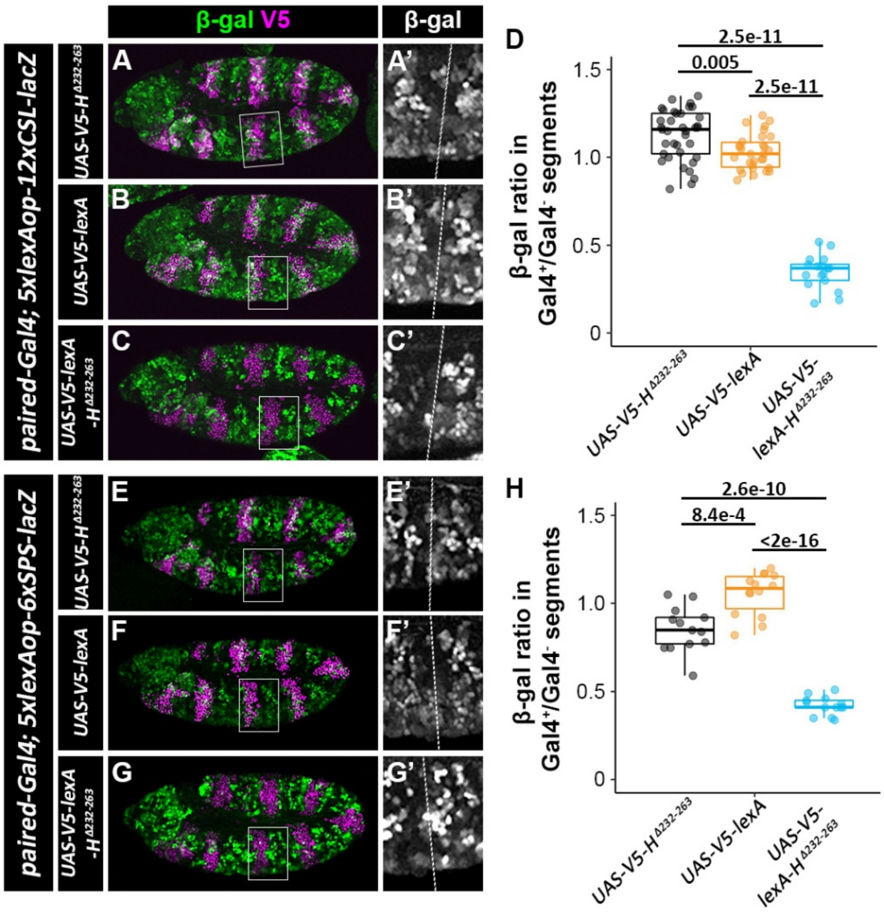


**Figure S6. lexA-H $^{\Delta232\text{-}263}$ fusion protein, but not lexA or H $^{\Delta232\text{-}263}$ alone, represses the lacZ reporters driven by lexA operator and CSL/SPS binding sites. A-C**. Stage 11 embryos of *paired-Gal4;5xlexAop-12xCSL-lacZ* with either *UAS-V5-Hairless*$^{\Delta232\text{-}263}$(A), *UAS-V5-lexA* (B) or *UAS-V5-lexA-Hairless*$^{\Delta232\text{-}263}$(C) immunostained with β-gal (green) and Hairless (magenta). A'-C'. Close-up views of β-gal intensity in black and white are shown in insets from A-C with the *paired-Gal4*-positive parasegment on the left and the *paired-Gal4*-negative parasegment on the right. **D**. Quantification of ratios of β-gal of *paired-Gal4*-positive over *paired-Gal4*-negative parasegments in *paired-Gal4;5xlexAop-12xCSL-lacZ* flies with indicated UAS construct. Each dot represents the average measurement from an individual embryo. Box plots show the median, interquartile range, and 1.5 times interquartile range. One-way ANOVA with post-hoc Tukey HSD was used to test significance. **E-G**. Stage 11 embryos of *paired-Gal4;5xlexAop-6xSPS-lacZ* with either *UAS-V5-Hairless*$^{\Delta232\text{-}263}$(E), *UAS-V5-lexA* (F) or *UAS-V5-lexA-Hairless*$^{\Delta232\text{-}263}$(G) immunostained with β-gal (green)

and Hairless (magenta). E'-G'. Close-up views of β-gal intensity in black and white are shown in insets from E-G. **H**. Quantification of ratios of β-gal of *paired-Gal4*-positive over *paired-Gal4*-negative parasegments in *paired-Gal4;5xlexAop-6xSPS-lacZ* flies with indicated UAS construct. Each dot represents the average measurement from an individual embryo. Box plots show the median, interquartile range, and 1.5 times interquartile range. One-way ANOVA with post-hoc Dunnett's T3 was used to test significance.
