## Supplemental Tables-Kuang et al for "Enhancers with cooperative Notch binding sites are more resistant to regulation by the Hairless co-repressor"

**Supplemental Tables for Kuang et al:**

**Supplementary Table 1**. EMSA probes.

| Probe name | Sequence |
|---|---|
| 2xCSL | CGAACGTGGGAAACCTAGGCTAGAGGCACCGTGGGAAACTAGTGCGGGCGTGGCT |
| 1xSPS | GCTACGTGGGAAAGGAGCAAACTGCGTTTCCCACGTTCGTAGTGCGGGCGTGGCT |
| 2xCSLmut | CGAACGAGGCAAACCTAGGCTAGAGGCACCGAGGCAAACTAGTGCGGGCGTGGCT |
| 1xSPSmut | |
| 5’IRDye-700 complementary_oligo | AGCCACGCCCGCACT |
| 5’IRDye-800 complementary_oligo | AGCCACGCCCGCACT |

**Supplementary Table 2**. Sequences used for molecular cloning.

| Enhancer | Restriction Site | Sequence |
|---|---|---|
| 2xCSL | EcoRI/BglII | GAATTCGCCCTGCGAACGTGGGAAACCTAGGCTAGAGGCACCGTGGG AAACTGCCTAGATCT |
| 4xCSL | EcoRI/BglII | GAATTCGCCCTGCGAACGTGGGAAACCTAGGCTAGAGGCACCGTGGG AAACTGCCTGCCCTGCGAACCGTGGGAAACCTAGGCTAGAGGCACCG TGGGAAACTGCCTAGATCT |
| 8xCSL | EcoRI/Acc65I | GAATTCGCCCTGCGAACGTGGGAAACCTAGGCTAGAGGCACCGTGGG AAACTGCCTGCCCTGCGAACCGTGGGAAACCTAGGCTAGAGGCACCG TGGGAAACTGCCTAGATCTGCCCTGCGAACGTGGGAAACCTAGGCTA GAGGCACCGTGGGAAACTGCCTGCCCTGCGAACCGTGGGAAACCTAG GCTAGAGGCACCGTGGGAAACTGCCTGGTACC |
| 12xCSL | EcoRI/BglII | GAATTCGCCCTGCGAACGTGGGAAACCTAGGCTAGAGGCACCGTGGG AAACTGCCTGCCCTGCGAACGTGGGAAACCTAGGCTAGAGGCACCGT GGGAAACTGCCTGCCCTGCGAACGTGGGAAACCTAGGCTAGAGGCAC CGTGGGAAACTGCCTGCCCTGCGAACGTGGGAAACCTAGGCTAGAGG CACCGTGGGAAACTGCCTGCCCTGCGAACGTGGGAAACCTAGGCTAG AGGCACCGTGGGAAACTGCCTGCCCTGCGAACGTGGGAAACCTAGGC TAGAGGCACCGTGGGAAACTGCCTAGATCT |
| 1xSPS | EcoRI/BglII | GAATTCAGCTACGTGGGAAAGGAGCAAACTGCGTTTCCCACGTTCGC AGGGCAGATCT |
| 2xSPS | EcoRI/BglII | GAATTCAGCTACGTGGGAAAGGAGCAAACTGCGTTTCCCACGTTCGC AGGGCAGCTACGTGGGAAAGGAGCAAACTGCGTTTCCCACGTTCGCA GGGCAGATCT |
| 4xSPS | EcoRI/Acc65I | GAATTCAGCTACGTGGGAAAGGAGCAAACTGCGTTTCCCACGTTCGC AGGGCAGCTACGTGGGAAAGGAGCAAACTGCGTTTCCCACGTTCGCA GGGCAGATCTAGCTACGTGGGAAAGGAGCAAACTGCGTTTCCCACGT TCGCAGGGCAGCTACGTGGGAAAGGAGCAAACTGCGTTTCCCACGTT CGCAGGGCGGTACC |
| 6xSPS | EcoRI/BglII | GAATTCAGCTACGTGGGAAAGGAGCAAACTGCGTTTCCCACGTTCGC AGGGCAGCTACGTGGGAAAGGAGCAAACTGCGTTTCCCACGTTCGCA GGGCAGCTACGTGGGAAAGGAGCAAACTGCGTTTCCCACGTTCGCAG GGCAGCTACGTGGGAAAGGAGCAAACTGCGTTTCCCACGTTCGCAGG GCAGCTACGTGGGAAAGGAGCAAACTGCGTTTCCCACGTTCGCAGGG CAGCTACGTGGGAAAGGAGCAAACTGCGTTTCCCACGTTCGCAGG GCAGATCT |
| 12xCSLmut | EcoRI/BglII | GAATTCGCCCTGCGAACGAGGCAAACCTAGGCTAGAGGCACCGAGGC AAACTGCCTGCCCTGCGAACGAGGCAAACCTAGGCTAGAGGCACCGA GGCAAACTGCCTGCCCTGCGAACGAGGCAAACCTAGGCTAGAGGCAC CGAGGCAAACTGCCTGCCCTGCGAACGAGGCAAACCTAGGCTAGAGG CACCGAGGCAAACTGCCTGCCCTGCGAACGAGGCAAACCTAGGCTAG AGGCACCGAGGCAAACTGCCTGCCCTGCGAACGAGGCAAACCTAGGC TAGAGGCACCGAGGCAAACTGCCTAGATCT |

| | | |
|---|---|---|
| 6xSPSmut | EcoRI/BglII | GAATTCAGCTACGAGGCAAAGGAGCAAACTGCGTTTGCCTCGTTCGC<br>AGGGCAGCTACGAGGCAAAGGAGCAAACTGCGTTTGCCTCGTTCGCA<br>GGGCAGCTACGAGGCAAAGGAGCAAACTGCGTTTGCCTCGTTCGCAG<br>GGCAGCTACGAGGCAAAGGAGCAAACTGCGTTTGCCTCGTTCGCAGG<br>GCAGCTACGAGGCAAAGGAGCAAACTGCGTTTGCCTCGTTCGCAGGG<br>CAGCTACGAGGCAAAGGAGCAAACTGCGTTTGCCTCGTTCGCAG<br>GGCAGATCT |
| 5xlexAop | HindIII/EcoRI | AAGCTTTCTGTATATATATACAGACGCAGTTTGCTCCTCTGTATATA<br>TATACAGTAGCTGCCCTGCGATCTGTATATATATACAGACGCAGTTT<br>GCTCCTCTGTATATATATACAGTAGCTGCCCTGCGATCTGTATATAT<br>ATACAGCCTAGGGTAGCATGCGTAACCGGTGTAGAATTC |
| LexA-DBD | NdeI/BglII | CATATGCCACCCAAGAAGAAGCGAAAAGTAGAAGATCCAATGAAGGC<br>TCTCACGGCCCGACAACAGGAAGTTTTTGATTTGATACGGGATCATA<br>TATCCCAAACGGGTATGCCTCCGACCCGCGCAGAGATAGCACAGCGA<br>CTGGGCTTTCGATCGCCTAACGCCGCGGAGGAGCACTTGAAGGCACT<br>GGCCCGCAAGGGTGTCATTGAAATCGTGTCCGGTGCGAGCCGCGGAA<br>TCCGGCTGTTGCAGGAGGAGGAAGAGGGCCTGCCACTGGTGGGACGC<br>GTGGCCGCTGGCGAGCCGCTGCTGGCCCAGCAACACATAGAGGGACA<br>CTATCAGGTGGACCCCTCCTTGTTTAAGCCAAATGCTGATTTCCTGT<br>TGCGGGTGTCGGGAATGTCCATGAAGGACATCGGTATTATGGATGGT<br>GACCTCCTCGCCGTCCATAAGACACAGGATGTCAGGAACGGCCAGGT<br>AGTCGTTGCCAGGATAGACGATGAGGTCACTGTGAAACGTCTCAAGA<br>AGCAAGGCAATAAGGTCGAGCTGCTGCCGGAGAATAGCGAGTTCAAG<br>CCGATCGTGGTGGATCTGCGACAGCAGTCCTTTACTATCGAGGGCTT<br>GGCCGTGGGTGTGATCCGCAACGGAGATTGGCTGGGCTCCGGCTCAG<br>ATCT |
| HairlessΔ23<br>2-263 | BglII/KpnI | AGATCTGATGGCCCTGCTTAATGACGTCACAAGCGTAGCAGAGTGCA<br>ACAGACAGACAACAATGACCGATGAGCATAAAAGTAACATTAACAGT<br>AACAGCAGTCACTCCAGCAACAACAACAACAACGGCAGCAGCAGCAA<br>TAACGACAACAACAGCAACGACGACGCAGCAAGTAGCAGCAACAGCA<br>AAAACAACAACACCAGCAACGAGAGCAGCCACAGCAACAACAATACT<br>AGTAGCATAATTGCAGAGGCGGCCGCCAAGTTTCTACTGAAAAATGG<br>CCTAAACGGCAGTAGCAGCACCAGCTACCCCCCTCTGCCACCGCCTC<br>TGCCCGCCAACTTAAGCAGGACGACCACGCCCACGACAACGACAACG<br>CCCTCATCCTCCAGCTCCACCGCCTCAAATGGCTTTTTGCCGCATGC<br>CAAGACGCCCAAAAGTAGTAGCATTATGGCTGCGTCCGCCGCAGTGG<br>CAGCCAGCGTCGTTGGAGCTACTGCGTCCAAGCCCACCATCGATGTC<br>CTGGGGGGCGTCCTGGACTACAGTTCCTTGGGCGGAGCTGCAACAGG<br>CTCACTGCCCACCACTGCAGTAGTAGCGGCGGCAGCGGGAACAGCGA<br>AGATCGGCAAGGGAAGCAACTCCGGCGGAAGCTTTGATATGGGCAGG<br>ACACCAATATCGACGCACGGCAACAACAGCTGGGGCGGCTACAAGAC<br>CTTCCGCCCTCCATCGGCGGCCACCTCCGCAACTGTGACCCCAACGT<br>CGGCGGTGACCACAGCGTATCCAAAGAACGAGAATTCCACATCGCTG<br>AGCTTTTCGGACGACAACAGCTCGATACAATCCTCTCCTTGGCAGCG<br>AGACCAGCCCTGGAAACAGTCCCGACCCAGGCGCGGCATATCTAAGG |

| | |
|---|---|
| | AGCTGTCGCTCTTCTTCCACCGCCCCAGGAACAGTACGCTTGGCCGA<br>GCTGCTCTCCGGACAGCCGCTCGCAAACGACGGCGCCCCCACGAGCC<br>GCTTACCACCAGCGAGGATCAGCAGCCCATTTTTGCGACGGCAATCA<br>AGGCGGAAAATGGAGACGATACTCTTAAAGCAGAAGCTGCAGAGGCC<br>GTTGAAATTGAAAATGTTGCTGTGGCGGACACAACCACAAATGAGAT<br>TAAAATTGAAAAACCGGACACGATCAAAGGCGAGGATGATGCTGAAC<br>GGCTCGAAAAGGAGCCGAAGAAGGCGGTTAGCGATGATAGCGAGTCA<br>AAAGAAGCATCGCCCGGTCAGCAAGTGGAACCACAACCAAAAGATGA<br>GACTGTTGATGTTGAGATGAAGATGAATACGAGCGAGGATGAGGAAC<br>CCATGACAGAGCTGCCCAGAATCACGAATGCCGTAAATGGTGATCTA<br>AACGGCGATCTAAAGGCGAGCATTGGGAAACCAAAATCCAAGCCGAA<br>GCCAAAAGCCAAGCTCAGCAGCATCATTCAGAAACTCATCGATAGCG<br>TACCAGCACGGCTTGAGCAAATGTCGAAGACATCAGCTGTGATCGCA<br>TCGACAACGACGTCTTCAGATCGCATTGGTGGCGGTCTAAGTCACGC<br>CTTGACGCACAAAGTTTCTCCACCCTCTTCTGCGACAGCAGCCGGAC<br>GACTAGTCGAGTACCACACCCAGCACGTGTCGCCCAGGAAAAGAATC<br>CTGCGCGAGTTCGAAAAGGTGTCGCTAGAGGACAACGGATGCGTAAA<br>CAACGGCAGCGGTGGAGCTAGTAGCGGTGGTGCTGGAGGAAAACGGA<br>GTCGAGCAAAGGGAACTTCGACATCGTCTCCGGCTGGCAAGGCGTCA<br>CCAATGAACTTGGCGCCACCCCAAGGAAAGCCAAGCCCCAGTCCCGG<br>CTCCAGCTCATCCAGCACTTCGCCAGCGACCTTGTCAACGCAGCCAA<br>CGCGGCTCAACAGCTCTTACAGTATCCACTCCCTGCTAGGTGGGAGC<br>AGTGGCAGCGGTAGCTCATCCTTCTCCTCCTCTGGCAAGAAGTGCGG<br>CGATCACCCGGCAGCTATTATCAGCAATGTGCACCATCCACAGCACT<br>CAATGTACCAACCCAGTTCCTCGAGCTATCCACGCGCCCTGCTCACC<br>TCGCCAAAGTCGCCCGATGTGAGTGGCAGCAATGGCGGGGGCGGAAA<br>ATCGCCCTCGCATACAGGAACCAAGAAGCGTTCGCCACCGTACTCGG<br>CGGGATCACCCGTAGACTATGGCCACTCCTTCTACAGGGATCCCTAT<br>GCGGGAGCAGGTCGTCCTTCCACATCGGGCTCAGCATCGCAGGACCT<br>GTCGCCACCGCGCTCTTCCCCAGCATCGCCAGCCACGACGCCGCGTA<br>CTGTGCCCAAAAAGACTGCATCGATCCGACGCGAGTTCGCTTCACCG<br>TCGGCCAGCAGCAGTAGCTGTCCCTCGCCCGGCGACCGGAGTGCATC<br>GCCCCCGGAACGGCGGCACATGCAGCAGCAGCCGCACCTACAGCGTA<br>GCTCGCCGCTGCACTACTATATGTACCCGCCACCGCCCCAGGTGAAC<br>GGGAACGGCTCGGCCGGAAGTCCGACCTCGGCGCCGCCCACGTCGAA<br>CAGCAGTGCAGCTGCAGTAGCGGCGGCAGCAGCGGCCGCAGCCGCAT<br>ACATTCCCTCGCCTTCGATATACAACCCGTACATATCCACACTGGCG<br>GCGTTGAGGCACAATCCGCTGTGGATGCACCACTATCAGACAGGAGC<br>GTCGCCCCTGCTGTCGCCACATCCACAACCCGGTGGCTCAGCGGCCG<br>CCGCTGCTGCAGCTGCTGCTGCGAGATTATCGCCCCAATCGGCCTAT<br>CACGCGTTCGCGTATAACGGAGTGGGAGCGGCTGTTGCCGCTGCAGC<br>AGCTGCGGCAGCCTTTGGACAACCGGCGCCCAGTCCCCACACGCATC<br>CGCACTTGGCCCATCCGCACCAGCATCCGCACCCGGCTGCACTGACC<br>ACCCACCACTCTCCCGCTCACCTGGCCACGCCAAAACTGACTGATAG<br>TAGTACCGACCAAATGTCTGCAACGTCCAGTCATCGCACAGCCTCCA |

| | |
|---|---|
| | CTTCGCCGAGCAGCTCGAGCGCATCGGCCTCCTCCTCGGCGGCCACT<br>TCGGGCGCCAGCTCCTCCGCAATGTTTCATACTAGTAGTCTAAGGAA<br>TGAACAAAGTTCAGACTTACCACTGAATCTGTCAAAGCACTGAGGTA<br>CC |
